# Abscission couples cell division to embryonic stem cell fate

**DOI:** 10.1101/798165

**Authors:** Agathe Chaigne, Celine Labouesse, Meghan Agnew, Edouard Hannezo, Kevin J Chalut, Ewa K Paluch

## Abstract

Cell fate transitions are key to development and homeostasis. It is thus essential to understand the cellular mechanisms controlling fate transitions. Cell division has been implicated in fate decisions in many stem cells, including neuronal and epithelial progenitors. In other stem cells, such as embryonic stem (ES) cells, the role of division remains unclear. Here we show that exit from naïve pluripotency in mouse ES cells generally occurs after a division. We further show that exit timing is strongly correlated between sister cells, which remain connected by cytoplasmic bridges long after division, and that bridge abscission progressively accelerates as cells exit naïve pluripotency. Finally, interfering with abscission impairs exit from naïve pluripotency. Altogether, our data indicate that a switch in the division machinery leading to faster abscission is crucial for pluripotency exit. Our study identifies abscission as a key step coupling cell division to fate transitions.

## Introduction

During embryonic development as well as in adult tissue homeostasis, cell fate transitions allow the generation of the diversity of cells comprising a functioning organism. For example, the zygotic cell is totipotent, as it can give rise to all the embryonic and extra-embryonic tissues, and embryonic development relies on a series of precisely controlled fate transitions. In the adult organism, stem cells, for example in the gut or the skin, produce the cell types needed for the maintenance of tissues like gut or lung (1). Understanding the cellular mechanisms underlying fate transitions is thus of fundamental importance for development and physiology. Cell division has been proposed to act as a switch during cellular fate transitions (2). A canonical example of mitotic control of cell fate is the first division of the C.elegans embryo, where cortical cues drive asymmetric spindle positioning, leading to a size asymmetry between daughter cells crucial for the specification of the antero-posterior axis (3). In most oocytes, size asymmetry during meiosis is essential to ensure that the fertilized oocyte retains the reserves essential for further embryo development, while the tiny polar body generated will degenerate (4). In Drosophila and C. elegans neuroblasts, asymmetries in daughter cell size upon division have been proposed to control cell fate (5, 6). Furthermore, in Drosophila and the zebrafish neural systems, as well as in Drosophila intestinal stem cells, the asymmetric segregation of Sara endosomes during cell division controls fate by modulating Notch signaling (7–10).

During early mammalian embryonic development, division asymmetries have been proposed to lead to acquisition of distinct fates by the two daughter cells (11). *In vitro*, cell division has been linked to cell fate choice in human ES cells: for example, when human ES cells submitted to primitive streak inducing signals divide, the two daughters cells often adopt different fates with one being resistant to differentiation (12). Moreover, artificially induced asymmetric divisions have been connected to mouse ES cell fate change: asymmetric division triggered by local application of beads coated with the signaling molecule Wnt3a, leads to the daughter cell distal from the Wnt signal expressing differentiation markers shortly after division (13). However, a number of studies suggest that in the absence of such external cues, lineage priming during exit from naïve pluripotency occurs in G1 phase (14–16). Consistent with this view, overall inhibition of cell division delays the downregulation of pluripotency genes in mouse ES cells exiting naïve pluripotency, though the underlying mechanisms remain unclear (15). Whether blocking cell division affects exit from naïve pluripotency functionally has not been tested. Altogether, the role of cell division for cell fate decisions in ES cells remains poorly understood.

Here we investigate the role of cell division in exit from naïve pluripotency using mouse ES cells as a model system. Using single-cell tracking, we show that exit from naïve pluripotency generally occurs after cell division, but that cells appear to need an entire cell cycle and a division for efficient exit. We then investigate how cell division affects the dissolution of the ES cell state. We show that dividing ES cells display strong size asymmetries between daughter cells, due to inhomogeneous distribution of E-cadherin in 3D colonies between the centre and the periphery. However, these asymmetries do not affect pluripotency exit dynamics. Instead, we show that sister cells display remarkably correlated naïve pluripotency exit timings, prompting us to test whether they remain connected even after division. Indeed, we find that abscission, the last stage of cell division when sister cells are physically separated, is slow in naïve ES cells, which remain connected by cytoplasmic bridges for a long time after division. Interestingly, abscission duration strongly decreases after naïve pluripotency exit is triggered. Finally, we demonstrate that interfering with abscission impairs naïve pluripotency exit. Altogether, we unveil a rewiring of the division machinery, leading to faster sister cell abscission, as a key step in exit from naïve pluripotency.

## Results

### ES cells exit pluripotency after mitosis

To investigate the role of cell division in exit from naïve pluripotency, we first tested the effect of inhibiting cell division altogether. To do so, we used ES cells expressing a short half-life naïve pluripotency reporter Rex1-GFPd2 (17), since the downregulation of Rex1-GFP correlates with exit from naïve pluripotency (17, 18). Cells were cultured in N2B27 medium supplemented with the MEK inhibitor PD0325901, the GSK3 inhibitor CHIRON, and Leukemia inhibitory factor (2i-LIF culture medium), and exit from naïve pluripotency was initiated by placing cells in N2B27 medium alone (exiting medium) (19). We blocked cell division with the CDK1 inhibitor RO-3306 and monitored Rex1-GFPd2 (hereafter Rex1-GFP) intensity after placing the cells in exiting media. While control cells showed a clear reduction of Rex1-GFP intensity 40 h after inhibitors removal, consistent with previous reports (17, 18), cells that did not undergo cell division maintained higher levels of Rex1-GFP (Figure S1A,B). The efficiency of RO-3306 to block cell division was confirmed by comparing bulk proliferation of control and RO-3306-treated ES cells (Figure S1C). Furthermore, the RO-3306-treated cells were found to increase their size dramatically, as expected for cells that are blocked in G2 (Figure S1D,E). These data suggest that cell division is important for exit from naïve pluripotency.

To further test the importance of cell division, we asked how its timing relates to exit from naïve pluripotency. We used the onset of Rex1-GFP downregulation as a readout of pluripotency exit timing. We first verified the dynamics of Rex1 downregulation at the population level. We observed that after 25-40h of exit, all cells had downregulated Rex1-GFP (Figure S1F, Supplementary Movies 1,2), consistent with previous reports (17). Furthermore, 24 hours after induction of pluripotency exit, the cells had downregulated key genes of the naïve pluripotency network (*Rex1, Klf2, Nanog, Klf4*) and upregulated genes typical of exit from pluripotency (*Fgf5, Otx2*) (Figure S1G). We then followed individual cells and their progeny to explore the correlation between cell division and Rex1-GFP downregulation (Figure 1A-D). The timing of Rex1-downregulation was determined automatically, as the time of the first inflexion of the curve in a sigmoidal fit to the time-course of Rex1-GFP intensity. Cell division appeared to correlate with exit from naïve pluripotency (Figure 1A, Supplementary Movie 1). Interestingly, some of the cells did not downregulate Rex1-GFP after the first division but they did so after undergoing a second division (Figure 1C,D, Supplementary Movie 2). Taken together, we found that the time of pluripotency exit strongly correlated with the time of the latest division (Figure 1E). Finally, we confirmed that the correlation between time of exit and time of division was very unlikely to be due to chance (Figure S1H and Methods). Altogether, these results show that the timing of exit from naïve pluripotency in ES cells correlates with cell division.

**Fig. 1:**
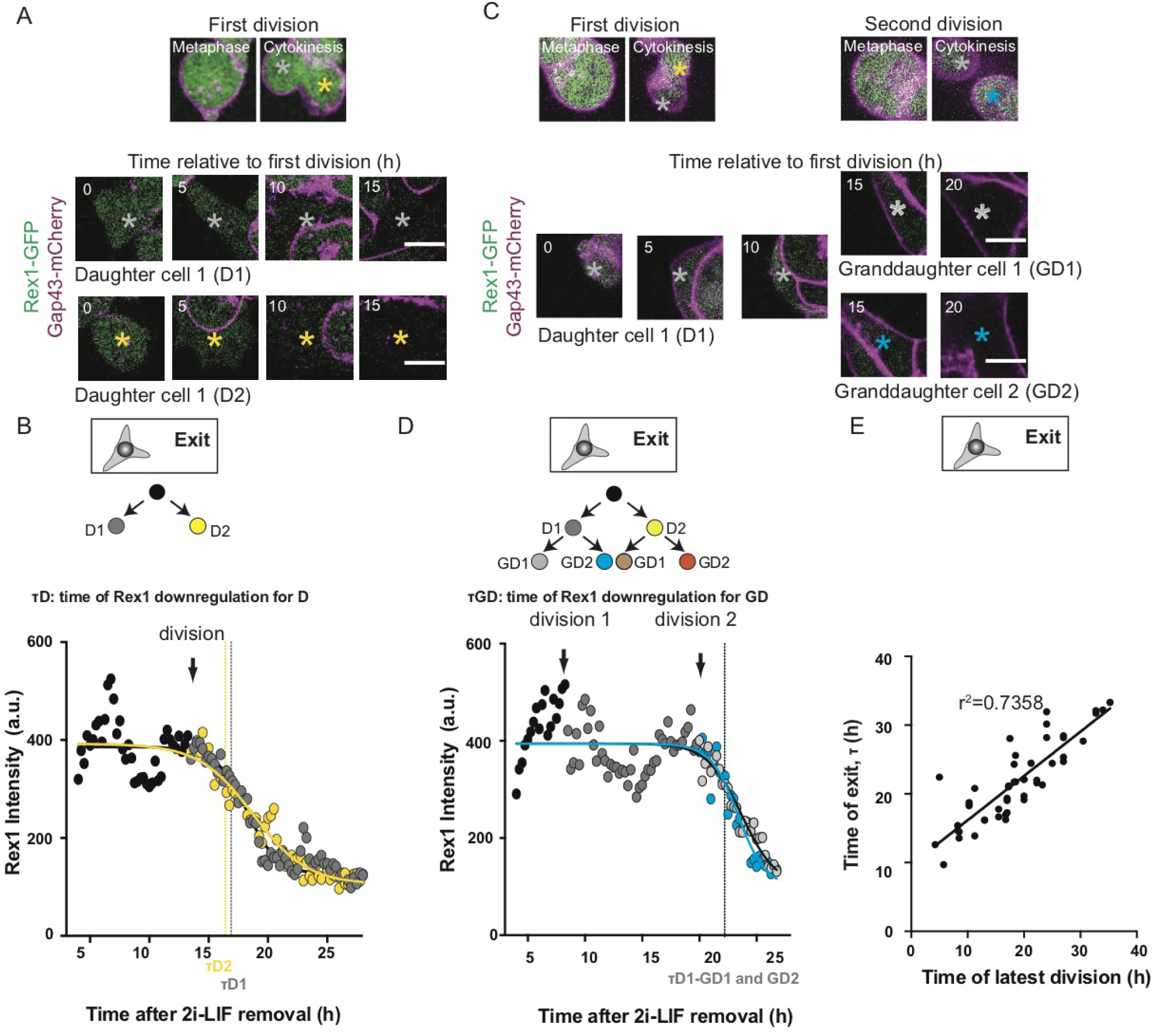
ES cells exit naïve pluripotency after mitosis. A) Representative example of an ES cell expressing Rex1-GFP (green) and Gap43-mCherry (magenta) undergoing one division before exiting pluripotency. Top: images of the cell division. Bottom: time-lapse of the two daughter cells highlighted in images at the top (Grey star: Daughter 1; yellow star: Daughter 2). One picture is shown every 1 hour, 0h: end of cytokinesis. A single Z-plane around the center of the cell at each time point is shown. Scale bar, 10 *µ*m. B) Plot showing Rex1-GFP mean intensity for the cells pictured in (A), as a function of time. 0h: time of 2i-LIF removal. Black dots: Mother cell. Grey dots: Daughter 1; yellow dots: Daughter 2. Lines are sigmoidal decay fits; the time of Rex1 downregulation (*τ*) is defined as the first inflexion of the curve (see Methods). The black arrow highlights the time of cell division. C) Representative example of ES cells expressing Rex1-GFP (green) and Gap43-mCherry (magenta) undergoing two divisions before exiting pluripotency. Top: images of the two cell divisions. Bottom: time-lapse of the daughter and grand-daughter cells highlighted in images at the top (Grey star: Daughter 1, Grand-daughter 1; blue star: Daughter 1, Grand-daughter 2). One picture is shown every 1 hour, 0h: end of the first cytokinesis. A single Z-plane around the center of the cell at each time point is shown. Scale bar, 10 *µ*m. D) Plot showing Rex1-GFP mean intensity for the cells pictured in (C) as a function of time. 0h: time of 2i-LIF removal. Black dots: Mother cell; grey dots: Daughter 1, Grand-daughter 1; blue dots: Daughter 1, Grand-daughter 2. Lines are sigmoidal decay fits; the time of Rex1 downregulation (*τ*) is defined as the first inflexion of the curve (see Methods). The black arrows highlight the times of the divisions. E) Scatter plot representing the time of Rex1-GFP downregulation (*τ*) as a readout of the time of naïve pluripotency exit, as a function of the time of the latest division. The latest division is determined as the division that happens before or up to 2.5h after (to account for experimental uncertainties in determining *τ*) the time of Rex1-GFP downregulation. 0h: time of 2i-LIF removal.

### ES cells go through a whole cell cycle and a division before exiting pluripotency

Since we observed that exit from pluripotency occurred shortly after a cell division, we hypothesized that inducing pluripotency exit in cells about to enter mitosis could result in faster exit. Thus, cells sorted at the end of the cell cycle should be the first ones to exit naïve pluripotency. To test this hypothesis, we used Fucci2a ES cells (20), which express two different fluorescent markers in different phases of the cell cycle, and sorted cells in distinct cell cycle phases. We then cultured cells in exiting media for 26h and assessed the effectiveness of naïve pluripotency exit using a clonogenicity assay (Figure 2A). In this assay, cells that have been cultured in exiting medium for a determined period of time are placed back in 2i-LIF, where only naïve pluripotent cells can survive; a low number of cells surviving in the assay is thus a readout of efficient pluripotency exit (Figure 2A, (18)). Surprisingly, we found that cells placed in exiting media at the exit from mitosis or while in G1 phase, exited pluripotency faster than control cells or cells synchronized in S/G2 phase, which are about to enter mitosis (Figure 2B).

**Fig. 2:**
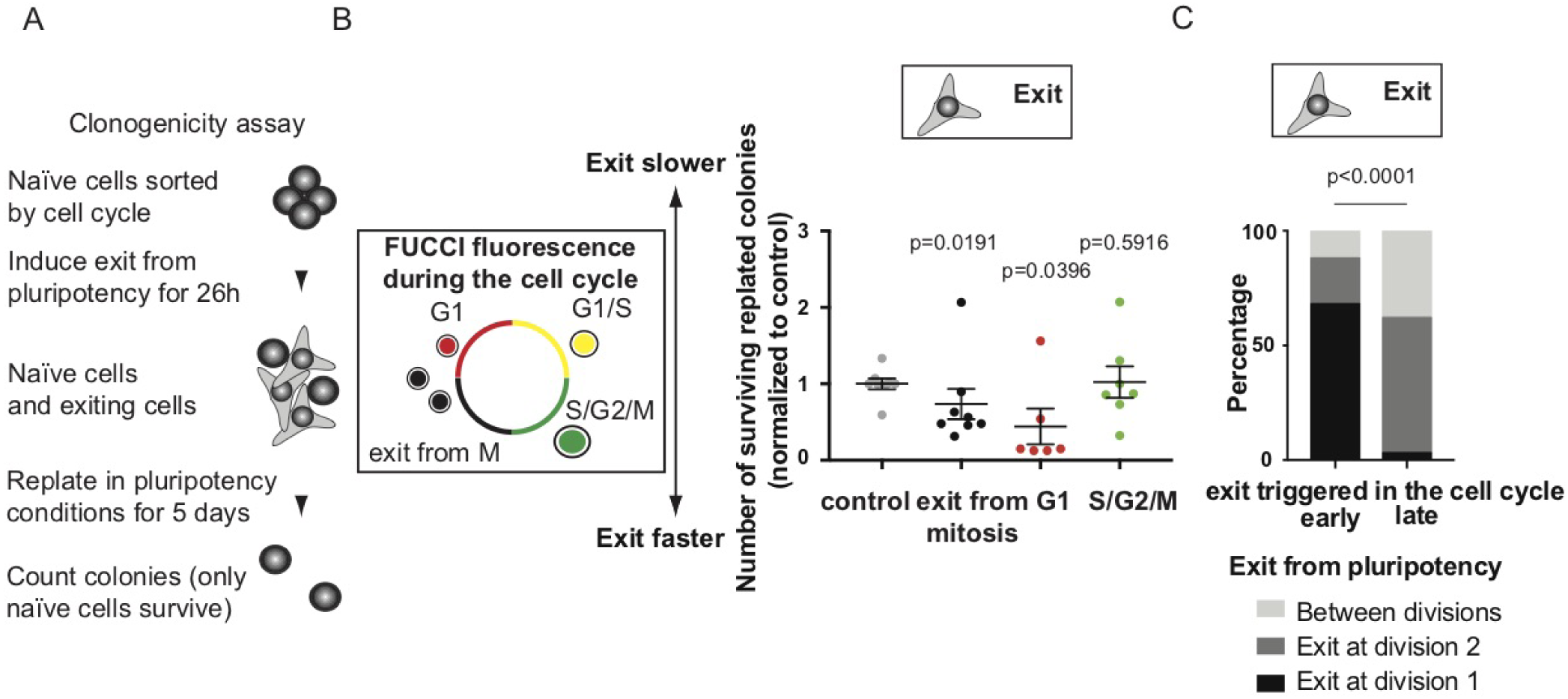
ES cells go through an entire cell cycle before exiting naïve pluripotency. A) Schematic of clonogenicity analysis assay (see Methods). B) Left panel: the Fucci system was used to sort cells by FACS. Cells in G1 express mCherry-Cdt1, cells at the G1/S transition start expressing mVenus-hGeminin; cells at the G1/S transition are double positive and cells exiting mitosis are double negative. Right panel: dot plot representing the number of ES cell colonies surviving in a clonogenicity assay performed on cells synchronized in different phases of the cell cycle. The control is the ungated population. The mean and standard error of the mean are shown. N=8. C) Contingency plot showing the percentage of cells downregulating Rex1-GFP around the time of the first division (± 4h, black), around the time of the second division (± 4h, dark grey), or in between two divisions (light grey) for cells where pluripotency exit is triggered early in the cell cycle (the first division happens more than 12h after 2i-LIF removal, left) or late in the cell cycle (the first division happens less than 12h after 2i-LIF removal, right). N=3, n=200.

To confirm these results at the single cell level, we sorted wild-type ES cells by size, as small cells largely correspond to cells that just exited mitosis or are in G1 phase (Figure S2A). We then performed single-cell RNA sequencing on the small (“early cell cycle”) cells, and on the unsorted population (“ungated”) after 6 hours in exiting media, in order to capture the first transcriptional changes of pluripotency exit. We first verified that the sorting conserved the cell cycle structure of the population, and observed that 6h after sorting and exit trigger, the majority of the initial “early cell cycle” population were in G1 or S phase, whereas the ungated population comprised mostly S phase and G2/M cells (Figure S2B). We then compared the expression levels of “early cell cycle” cells to the control population and found that after 6 hours in exiting media, the cells for which exit was triggered early in the cell cycle displayed overall stronger downregulation of pluripotency genes than the ungated population (Figure S1C). Furthermore, a cluster analysis separating the cells based on expression levels of two of the earliest genes down-regulated during pluripotency exit, *Tfcp2l1* and *Tbx3* (17), indicated that “early cell cycle” cells displayed a stronger downregulation of these early genes compared to the ungated population (Figure S1D). Taken together, these results suggest that cells exit pluripotency faster when exit is triggered in cells that just exited mitosis (Figures 2A,B and S2), yet mitosis appeared to be important for loss of naïve pluripotency (Figure 1).

To understand this paradox, we took a closer look at the correlation between cell division and Rex1-downregulation (Figure 1). We separated the cells into cells that divided shortly after 2i-LIF removal (less than 12h), which means 2i-LIF was removed late in the cell cycle, and cells that divided late after 2i-LIF removal, which means 2i-LIF was removed early in the cell cycle. Cells that were triggered to exit naïve pluripotency early in the cell cycle mostly downregulated Rex1-GFP after the first division, and cells triggered late in the cell cycle predominantly started downregulating Rex1-GFP after the second division (Figure 2C). Altogether, these results suggest that ES cells need to go through an entire cell cycle and a division before exiting from naïve pluripotency.

### ES cells present strong size asymmetries between daughter cells at cell division

We then explored how division affected exit from naïve pluripotency. Since asymmetric divisions, in particular in size, are important for fate specification in a number of stem cell types (3, 5, 6, 12), we asked if ES cells display asymmetries at cell division. We monitored cell divisions in ES cells stably expressing H2B-RFP to label DNA (21). We tested that the cell line could contribute to all the tissues by injecting it in a blastocyst, which gave rise to a viable mouse (Figure S3A). We acquired time lapses of H2B-GFP ES cells in 3D colonies labeled with CellMask™ to mark the membrane, and used a custom plugin that we previously developed (22) to measure cell volume throughout cell division (Figure S3B). Single isolated ES cells divided mostly symmetrically, as previously described (13). However, ES cells normally grow in 3D colonies. We found that when dividing in colonies, ES cells displayed significant asymmetries between daughter cell sizes (Figure 3A,B, Supplementary Movies 3,4). Cells dividing at the periphery of the colonies displayed stronger asymmetries than cell dividing inside colonies, and cells at the periphery of a colony with the spindle oriented perpendicularly to the colony border (“radial” orientation) displayed highest size asymmetries between daughter cells (Figure 3A,B). As a comparison, HeLa cells, heavily derived cancer cells with great variability in chromosome count which are thus not expected to control their size very precisely, divided much more symmetrically than mouse ES cells (Figure 3B and Figure S3C, Supplementary Movie 5).

**Fig. 3:**
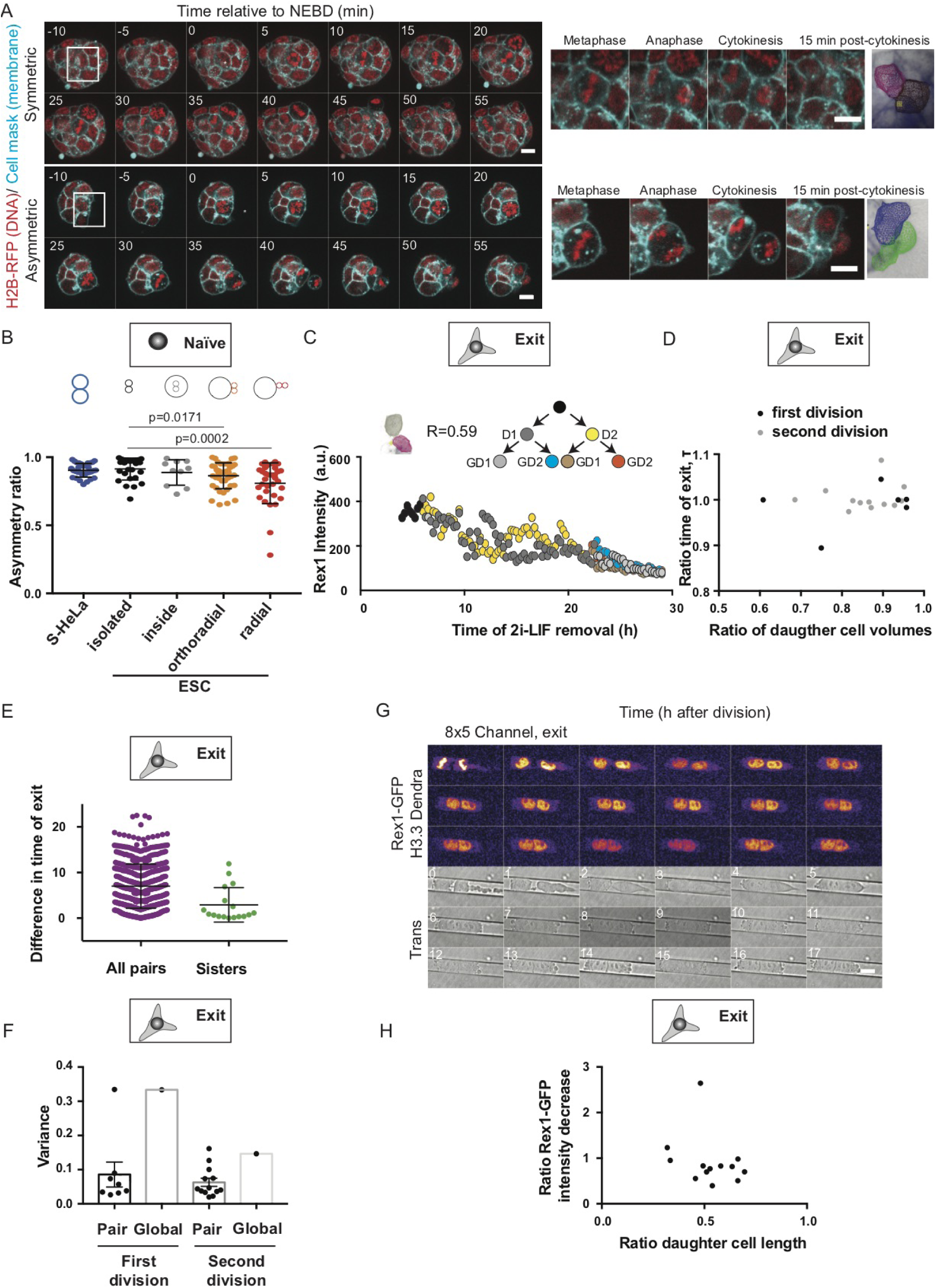
ES cells display strong size asymmetries at cell division, yet daughter cells are correlated in naïve pluripotency exit dynamics. A) Left: Representative time-lapses of colonies of naïve ES cells expressing H2B-RFP (red) and labeled with CellMask^*TM*^ far red (cyan) with one cell dividing at the center (top) and at the periphery of the colony (bottom); and close up images (right) of the regions highlighted with white boxes. Time in min; 0 min: time of nuclear envelope breakdown (NEBD). A single Z-plane is shown. Scale bars: 10 *µ*m. B) Dot plot representing the size asymmetry ratio between daughter cells for S-HeLa (blue), single ES cells (“isolated”, black), ES cells dividing at the center of a colony (“inside”, grey), and ES cells dividing at the periphery of a colony with the mitotic spindle oriented parallel (“orthoradial”, orange) or perpendicular (“radial”, red) to the colony border. The mean and standard deviation is plotted. N=3. C) Plot of Rex1-GFP mean intensity for ES cells exiting pluripotency after a very asymmetric division as a function of time. 0h: time of 2i-LIF removal. Inset: 3D-meshworks showing the volumes of the daughter cells and corresponding asymmetry ratio. D) Plot showing the ratio between the times of Rex1 downregulation for sister cells exiting shortly after the first (black) or second (grey) division as a function of the ratio of the volumes of the sister cells. 0h: time of 2i-LIF removal. N=3, n=18 pairs of sisters. E) Dot plot showing the difference in exit time for pairs of cells chosen at random (left, purple) and pairs of sister cells (right, green). The mean and standard deviation are plotted. N=3. F) Dot plot showing the variance of the intensity of the Rex1-GFP signal at every time point (see Methods) for cells exiting at the first division (left) or the second division (right), either compared to the global average of all cells (“global”) or between pairs of sister cells (“pair”). N=3. G) Time-lapse of an ES cell expressing H3.3-Dendra and Rex1-GFP (fire, upper panel) dividing in an 8*5 *µ*m cannel in N2B27. The transmitted light channel used for monitoring cell length is shown in the bottom panel. One picture is shown every 1 hour. 0 min corresponds to anaphase. One Z-plane is shown. Scale bar: 10 *µ*m. H) Plot showing the ratio of Rex1-GFP intensity decrease as a function of the ratio of daughter cell size 6h after cell division in the 8*5 *µ*m channels. N=6.

We then explored the mechanisms underlying asymmetric division in ES cells. We noted that cells at the periphery displayed significant shape instabilities during cell division (Figure S3D, Supplementary Movie 6), correlating with enhanced division size asymmetries (Figure S3E). Myosin-driven contractions have been shown to induce shape instabilities in dividing cells (23), and lead to asymmetric divisions in neuroblasts (6, 24). We found that myosin accumulated at the border of ES cell colonies (Figure S3F,G). However, myosin inhibition with low doses of Blebbistatin, which slowed down cytokinesis (Figure S3H) without preventing cell division, reduced cytokinetic shape instabilities but did not reduce division asymmetries (Figure S3I,J, Supplementary Movies 7). We then hypothesized that E-Cadherin, which controls cell-cell junctions and is thus absent from colony borders, may be responsible for division asymmetries. To test this, we plated cells on E-Cadherin-coated substrates, which lead to naïve ES cells spreading in 2D colonies (Figure S3K). Cells plated on E-Cadherin divided much more symmetrically than cells in 3D colonies (Figure S3K-M, Supplementary Movie 8). Enhanced division symmetry was not a mere effect of colony spreading, as cells at the periphery of ES cell colonies plated on laminin, which also adopt 2D morphologies (Figure S3K), displayed division asymmetries comparable to cells in 3D colonies (Figure S3N, Supplementary Movie 9). Altogether, these results suggest that inhomogeneity in E-cadherin distribution along the contour of cells dividing at the periphery of ES cell colonies leads to strong asymmetries in daughter cell size.

### Exit from pluripotency is insensitive to size asymmetries at cell division

We then asked if asymmetries in daughter cell size affect naïve pluripotency exit dynamics. We did not observe significant differences in Rex1-GFP intensity dynamics between the two daughter cells, even when the division was very asymmetric in size (Figure 3C,D). In fact, the timing of Rex1 downregulation was very correlated between sister cells (Figure 3E) and the variance of Rex1-GFP levels was very low between sisters (Figure 3E,F). These data indicate that sister cells undergo naïve pluripotency exit in a highly correlated manner, and that size asymmetries at cell division do not influence the timing of pluripotency exit. To directly test this, we induced strongly asymmetric divisions by confining naïve ES cells in narrow microchannels, as confinement has been shown to induce asymmetries at cell division in other cell types (25, 26). Confinement reliably induced division asymmetries in ES cells (Figure S4A,B, Supplementary Movie 10). We found that in the hours following cell division in microchannels, Rex1-GFP levels displayed similar levels in the two daughter cells (Figure 3G, Figure S4C), and no correlation was observed between the size ratio of the daughter cells and the ratio of Rex1-GFP intensity decrease 6h after cell division in the two daughter cells (Figure 3H). In conclusion, size asymmetries between daughter cells at cell division do not appear to influence the timing of exit from naïve pluripotency.

### Sister cells remain connected after division in ES cells

The strong correlation in Rex1 downregulation dynamics between daughter, and in some cases grand-daughter cells (Figure 3E,F) led us to ask whether daughter cells might remain connected after division in ES cells and during early stages of exit from naïve pluripotency. We thus imaged microtubules and observed that naïve ES cell colonies displayed a high number of tubulin bridges, remnants of mitotic spindles, still connecting daughter cells (Figure 4A). We then asked if sister cells connected by a bridge could still exchange cytoplasmic material. We expressed cytoplasmic GFP and used photo-bleaching to abruptly decrease cytoplasmic intensity. We found that photo-bleaching cytoplasmic fluorescence in one sister cell lead to a decrease in fluorescence intensity in the other sister, but not in a nearby unconnected cell positioned at a similar distance (Figure 4B-D, Supplementary Movie 11). Taken together, these results suggest that abscission is slow in ES cells and that sister ES cells remain physically connected after cell division and exchange cytoplasmic material.

**Fig. 4:**
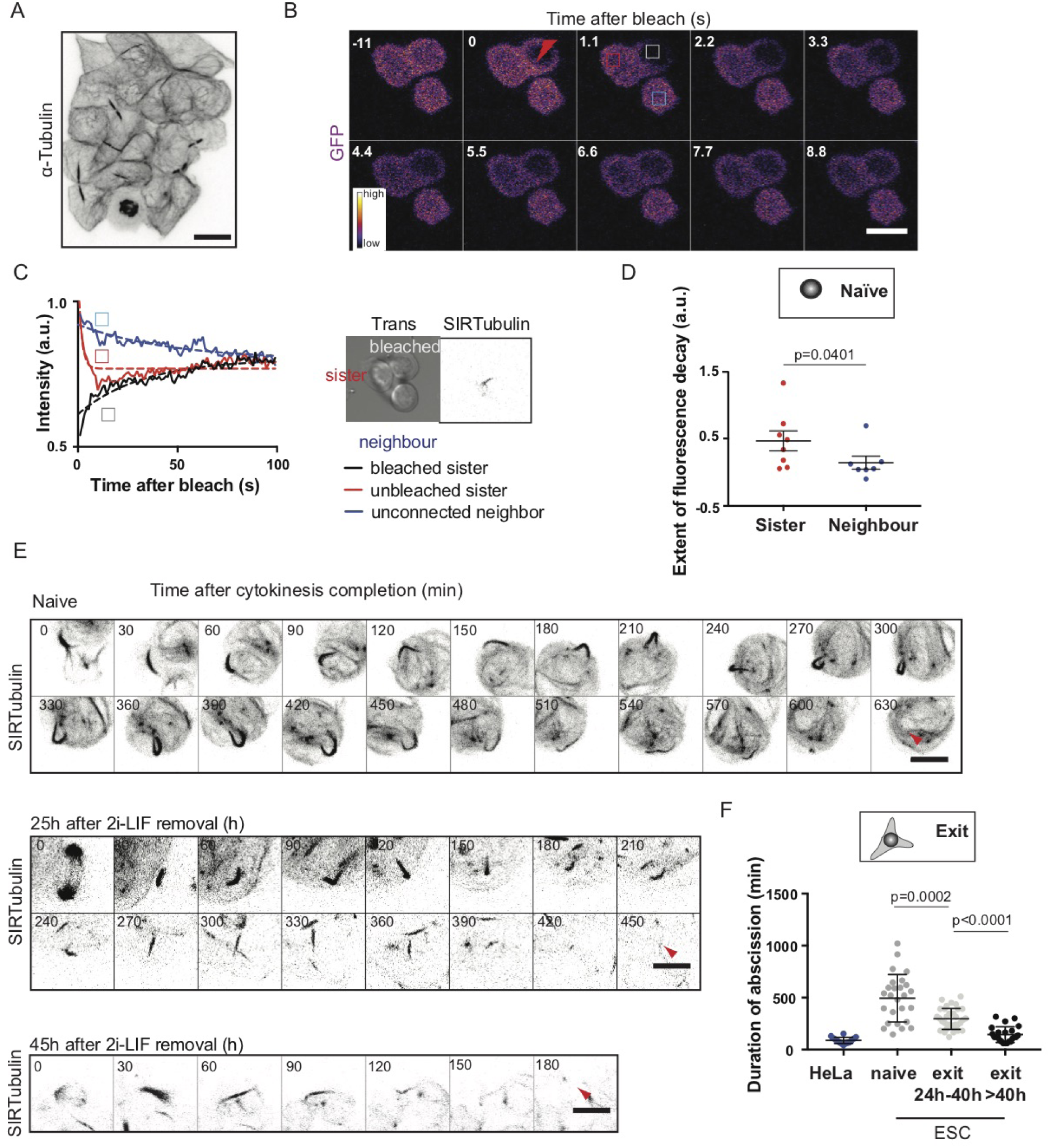
Naïve ES cells display long abscission times. A) Representative confocal image of a naïve ES colony stained for *α*-Tubulin (black, inverted contrast). A maximum Z-projection of shown over the volume of the colony. Scale bar: 10 *µ*m. B) Representative time-lapse of a FRAP experiment in ES cells expressing GFP, the two cells at the top are sister cells. The intensity levels are displayed. Photobleaching is performed at 0s in the sister cell on the right (red thunderbolt). The intensity is then measured in the bleached cell (dark grey box), the sister cell (red box) and an unconnected neighbor (blue box). One Z-plane is shown, scale bar: 10 *µ*m. C) Left panel: Plot showing the GFP intensity of the cells depicted in (B) and fitting curves. Black: bleached cell, red: sister cell, blue: unconnected neighbor. Right panel: transmitted light image and fluorescent Z-projection (inverted contrast) of the 3 cells displayed in (B) and labeled with SIRTubulin prior to the FRAP experiment. The two top cells are connected by a tubulin bridge. D) Dot plot showing the extent of the decay of the EGFP signal for the sister cell of the ES cell where GFP was bleached (red) and for an unconnected neighboring cell at a similar distance (blue). The mean and standard error of the mean are plotted. N=3. E) Representative time-lapses of a colony of ES cells expressing H2B-RFP and labeled with SIR-Tubulin (black, inverted contrast, the maximum Z-projection across the colony is shown). One picture is shown every 30 min. 0 min: end of cytokinesis. Top: naïve cells. Middle: cells 25h after induction of pluripotency exit, bottom: cells 45h after induction of pluripotency exit. Red arrows: abscission. Scale bars: 10 *µ*m. F) Dot plot showing the duration of abscission for HeLa cells expressing Tubulin-GFP dividing on elongated line micropatterns to standardize cell shape (blue) and for naïve ES cells and cells exiting pluripotency labeled with SIR-Tubulin (grey and black dots). The mean and standard error of the mean are shown. N=3.

### Abscission duration decreases during exit from naïve pluripotency

Since sister cells remain physically coupled by tubulin bridges after division, and appear to exit naïve pluripotency with similar dynamics, we hypothesized that abscission, the last step of cell division during G1 phase, could be important for pluripotency exit. To characterize abscission dynamics during exit from naïve pluripotency, we acquired time-lapse movies of cells treated with low doses of SIR-Tubulin (27), a live marker of tubulin, and then measured the time of abscission after cytokinesis (Figure 4E,F, Supplementary Movies 12-14). We found that naïve ES cells maintained tubulin bridges for a much longer time compared to HeLa cells (8.2±3.8h in naïve ES cells versus 1.5±0.5h in HeLa cells, Figure 4E,F), further indicating that abscission takes a long time in ES cells. Interestingly, the duration of abscission significantly decreased during exit from naïve pluripotency (Figure 4E,F). To further explore changes in abscission during pluripotency exit, we immuno-stained tubulin and the midbody marker Citron Rho-interacting kinase (CRIK) in naïve ES cells and cells at various stages of exit (Hu et al., 2012). All bridges were found to display CRIK foci, but some CRIK foci were not associated with bridges, suggesting they may mark midbody remnants (Figure 5A). We found that bridge density gradually decreased (Figure 5A,B), while the number of midbody remnants increased (Figure S5A) during exit from naïve pluripotency. Together, these results indicate that abscission is slow in naïve ES cells, and that the duration of abscission decreases during pluripotency exit.

**Fig. 5:**
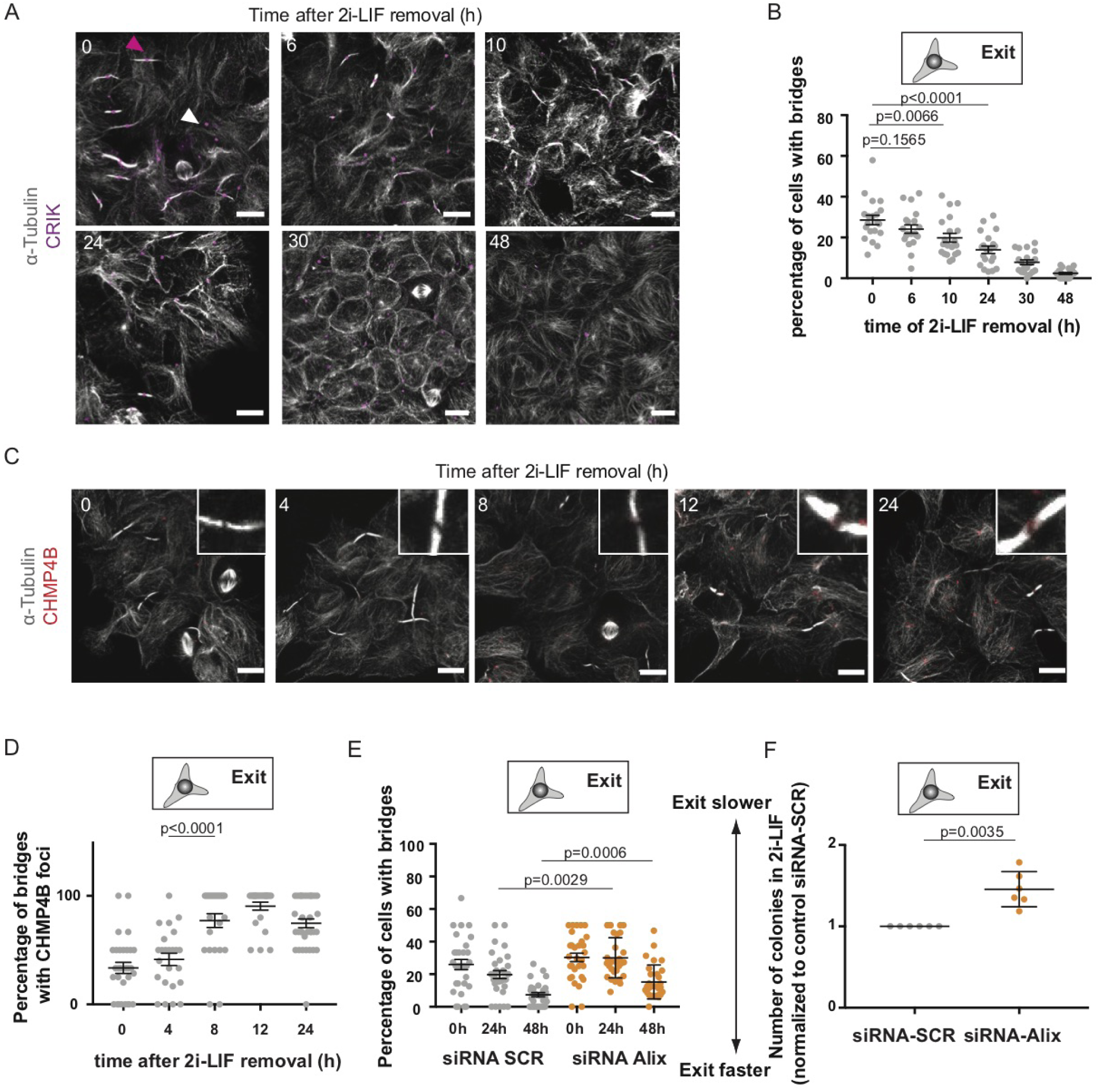
Abscission regulates exit from naïve pluripotency. A) Representative confocal images of ES cells at different stages of naïve pluripotency exit and stained for *α*-Tubulin (white) and CRIK (magenta). Pink arrowhead: example of a bridge with a CRIK spot; white arrowhead: example an isolated CRIK spot, suggesting a midbody remnant. Cells are cultured on laminin to facilitate the visualization of the bridges. Scale bars: 10 *µ*m. B) Dot plot showing the percentage of cells with bridges detected in H2B-RFP ES cells during exit from pluripotency on laminin. The mean and standard error of the mean are shown. N=2. C) Representative confocal images showing ES cells stained for *α*-Tubulin (white) and CHMP4B (red) during exit from pluripotency. Inset: zoom of a representative bridge. Scale bars: 10 10 *µ*m. D) Dot plot showing the percentage of bridges with a CHMP4B spot in the vicinity of the bridge center in ES colonies during exit from naïve pluripotency on laminin. The mean and standard error of the mean are shown. N=2. E) Dot plot showing the percentage of cells with bridges in ES colonies treated with siRNA Scrambled (grey) or Alix (orange) for 24h in 2i-LIF during exit from naïve pluripotency on laminin. The mean and standard error of the mean are shown. N=2. F) Dot plot representing the number of colonies surviving in a clonogenicity assay (see Figure 2A) for ES cells treated with siRNA Scrambled (SCR, grey) or Alix (orange) for 24h in 2i-LIF then placed in exiting media for 24h. The mean and standard deviation are shown. N=6.

We then asked whether the faster abscission dynamics during pluripotency exit could result from molecular changes in bridge composition. Notably, the expression levels of key known abscission regulators do not appear to change during exit from naïve pluripotency (Table S1, original data from (17, 28)). We thus probed changes in localization, focusing on the ECRT-III protein CHMP4B. CHMP4B performs the last step of abscission by polymerizing into circular filaments that are thought to cut the bridge (29, 30), and as such can be used as a readout of bridge maturity. We found that only 33% of the bridges in naïve ES cells displayed CHMP4B foci at their center (Figure 5C,D). However, the proportion of bridges with a CHMP4B spot at the center of the bridge steadily increased during pluripotency exit (Figure 5C,D), with a strong increase in proportion of bridges with CHMP4B between 4h and 8h after triggering exit from naïve pluripotency (Figure 5D). Together, these data suggest that the abscission machinery is remodeled during exit from pluripotency, leading to faster abscission after about 8 hours.

### Interfering with abscission impairs exit from naïve pluripotency

Finally, we asked whether interfering with abscission affects exit from naïve pluripotency. We depleted the ESCRT-III complex upstream activator ALIX, which allows to impair abscission (31, 32), without significantly affecting intercellular trafficking processes (33). In naïve ES cells, siRNA against ALIX did not affect the expression of key pluripotency markers but efficiently decreased ALIX expression (Figure S5B). ALIX depletion impaired the decrease in bridge density after induction of pluripotency exit, suggesting that it effectively targets abscission (Figure 5E, Figure S5C). We then performed a clonogenicity assay and found that ALIX depletion impaired exit from naïve pluripotency (Figure 5F). We confirmed these results by analyzing the expression levels of pluripotency genes 24h after triggering exit from naïve pluripotency in control and ALIX-depleted cells. We found that several key pluripotency genes, including *Nanog* and *Klf4*, maintained high expression levels in ALIX-depleted cells (Figure S5D). Finally, we verified that the effect of abscission on exit from pluripotency was not due to the specific culture conditions, so we repeated these experiments using an alternative pluripotency-promoting culture medium (Serum-LIF). We found that when ES cells exited naïve pluripotency from Serum-LIF, they also presented a decrease in bridge density (Figure S5E), an increase in mid-body density (Figure S5F), and ALIX siRNA also impaired exit from pluripotency (Figure S5G). Altogether, these results suggest that a quicker resorption of intercellular bridges after removal of pluripotency-promoting media promotes exit from naïve pluripotency.

## Discussion

Using a combination of functional assays and single cell tracking, we have shown that exit from naïve pluripotency, as assessed by the timing of Rex1 downregulation (17), occurs after cell division. When exit is induced early in the cell cycle, cells downregulate Rex1 after the first division, whereas when exit is induced later in the cell cycle, cells generally undergo two divisions before Rex1 downregulation (Figures 1 and 2). We conclude that cells need an entire cell cycle and a division to effectively exit pluripotency. This is consistent with a previous report showing that when exit is triggered in cells synchronised in G1 phase, pluripotency exit is initiated during the same cell cycle (15).

Previous studies have identified the G1 phase as a key stage for triggering pluripotency exit, as it allows for differentiation signals to rewire gene expression during DNA replication in S phase and because of direct control of pluripotency factors by G1 cyclins (15, 16). Here, we identify the last step of mitosis, abscission, which also happens in G1 phase (34), as a permissive cue for exit from pluripotency. We found that abscission is very slow in mouse ES cells, leading to cells remaining connected by cytoplasmic bridges for a long time after cell division, and that abscission progressively accelerates during pluripotency exit. This is in line with previous observations that enhanced midbody release, and thus enhanced abscission, accompany cell differentiation in a number of cell types (35). Here, we demonstrate that maintaining bridges impairs exit from naïve pluripotency (Figure 5). Interestingly, cells in the early mouse embryo have been shown to retain tubulin bridges throughout interphase from the 2-cell stage up to the blastocyst stage, and these interphase tubulin bridges have been proposed to act as a platform for E-Cadherin transport towards cell-cell junctions (36). E-Cadherin has been implicated in pluripotency maintenance (37), thus, long-lived tubulin bridges could help maintain pluripotency in ES cells by ensuring E-Cadherin targeting to cell-cell contacts.

It will be interesting to further explore how the abscission machinery is remodelled during pluripotency exit. The levels of the main known abscission regulators do not display strong changes during exit from naïve pluripotency (Table S1, (17, 28)), suggesting that accelerated abscission results from changes at the protein level, possibly via controlling the addressing of essential factors to the bridge. Our data suggest a role for the ESCRT-III protein CHMP4B, a key driver in the physical resolution of the bridge (for review see (38)), which is progressively recruited to the bridges after induction of pluripotency exit. Interestingly, in a recent study we identified a decrease in plasma membrane tension as a key regulator of exit from pluripotency (de Belly et al., *submitted*). Abscission efficiency has been linked to membrane tension, with high membrane tension acting as a negative regulator of abscission in HeLa cells by preventing the recruitment of ESCRT-III proteins (39). It is tempting to speculate that membrane tension decrease during pluripotency exit could also control the recruitment of the ESCRT machinery to the bridge and result in faster abscission.

Finally, we also observed that naïve mouse ES cells, and cells exiting pluripotency, display significant size asymmetries at cell division. Embryonic stem cells have been generally thought as dividing symmetrically in culture (40), unless provided with external cues (13). Here we show that ES cells often display high size asymmetries due to inhomogeneous distribution of E-Cadherin around ES cells in 3D colonies (Figure S3). As mammalian cells usually regulate their size by adjusting the length of their G1 phase (26), size heterogeneities between daughter cells could enhance variability in cell cycle length. Based on our observation that pluripotency exit requires an entire cell cycle, size heterogeneities could then directly lead to variability in pluripotency exit timings. As noise at the population level has been proposed to be important for cell fate decisions (41), it will be interesting to investigate if asymmetric cell divisions could contribute to generating noise in pluripotency exit dynamics.

In conclusion, our data uncover how changes in a key cell biology process, the separation of sister cells during abscission, act as a permissive cue for pluripotency exit. These results shed light on how modulating the dynamics of specific cell cycle processes can contribute to cell fate transitions.

## ACKNOWLEDGEMENTS

We would like to thank the entire Paluch and Baum lab at the MRC/LMCB and the Chalut lab at the SCI for discussions and feedback throughout the project, the microscopy platform in particular Andrew Vaughan and the flow cytometry platform in particular Stephanie Canning for their invaluable expertise and support. We would like to thank Jonathan Chubb (MRC/LMCB) and Carla Mulas (SCI, University of Cambridge) for providing technical expertise and advice. AC would like to think in particular Buzz Baum and Gautam Dey for helpful discussions.

## AUTHOR CONTRIBUTIONS

AC and EKP designed the research and wrote the manuscript; AC performed all the experiments and analyzed the data. CL performed the sc-RNAseq analysis. EH analyzed the Rex1GFP tracking. MA made the Dendra-H3.3 Rex1-GFP cell line. KJC and EKP supervised the project. All authors provided inputs on the project and the manuscript.

## FUNDING

This work was supported by the Medical Research Council UK (MRC programme award MC-UU-12018/5, AC, MA and EKP), the European Research Council (starting grant 311637-MorphoCorDiv to EKP), the Leverhulme Trust (Leverhulme Prize in Biological Sciences to EKP). KJC acknowledges support from the Royal Society (Royal Society Research Fellowship). AC acknowledges support from EMBO (ALTF 2015-563), the Wellcome Trust (201334/Z/16/Z), and the Fondation Bettencourt-Schueller (Prix jeune chercheur 2015).

## METHODS

### Data and materials availability

All data and materials used in the analysis are available upon request.

### Cell culture, live imaging and transfection

Mouse embryonic stem cells were routinely cultured as described in (Mulas et al., 2019) on 0.1% gelatin in PBS (unless otherwise stated) in N2B27+2i-LIF + penicillin and streptomycin, at a controlled density (1.5-3.0 104 cells/cm2) on Falcon flasks and passaged every other day using Accutase (Sigma-Aldrich, #A6964). They were kept in 37C incubators with 7% CO_2_. Cells were regularly tested for mycoplasma.

In this study, the cells used were: E14 wild type cells, E14 cells stably expressing H2B-RFP (21), E14 cells stably expressing Rex1-GFP and Gap43-mCherry (a kind gift from Carlas Mulas, Stem Cell Institute, University of Cambridge) (17), 3T3 cells expressing the Fucci2a system (20) (a kind gift from Ian James Jackson, the University of Edinburgh). The E14 cells stably expressing H2B-RFP (21) was tested for contribution to chimera by injection at the Francis Crick Institute mouse facility into a C57B6 blastocyst and gave rise to a viable mouse with good contribution of the H2B-RFP cells as assessed by the color of the mouse and dissection under a fluorescent lamp (Figure S3, the mouse is white despite a C57Bl6 background).

The culture medium was made in house, using DMEM/F-12, 1:1 mixture (Sigma-Aldrich, #D6421-6), Neurobasal medium (Life technologies #21103-049), 2.2 mM L-Glutamin, home-made N2 (see below), B27 (Life technologies #12587010), 3 *µ*M Chiron (Cambridge Bioscience #CAY13122), 1 *µ*M PD 0325901 (Sigma-Aldrich PZ0162), LIF (Merck Millipore # ESG1107), 50 mM *β*-Mercapto-ethanol, 12.5 ng.mL-1 Insulin zinc (Sigma-Aldrich #I9278). The 200 X home-made N2 was made using 0.791 mg.mL-1 Apotransferrin (Sigma-Aldrich #T1147), 1.688 mg.mL-1 Putrescine (Sigma-Aldrich #P5780), 3 *µ*M Sodium Selenite (Sigma-Aldrich #S5261), 2.08 *µ*g.mL-1 Progesterone (Sigma-Aldrich #P8783), 8.8% BSA. Exit from pluripotency was triggered by passaging the cells and seeding them in N2B27.

For colony imaging, the cells were typically plated on 35 mm Ibidi dishes (IBI Scientific, #81156) coated with gelatin (unless otherwise stated) the day before the experiment, and imaged on a Perkin Elmer Ultraview Vox spinning disc (Nikon Ti attached to a Yokogawa CSU-X1 spinning disc scan head) using a C9100-13 Hamamatsu EMCCD Camera. Samples were imaged using a 60X water objective (CFI Plan Apochromat with Zeiss Immersol W oil, Numerical Aperture 1.2). Typically, the samples were imaged overnight acquiring a Z-stack with ΔZ = 2 *µ*m every 5 minutes.

Transfections were performed using 5 *µ*g of plasmid and 6 *µ*L of Lipofectamin, incubated in 250 *µ*L OptiMEM for 5 minutes, then mixed and incubated at room temperature for 20 minutes, and added to cells passaged onto Ibidi dishes concomitantly. The medium was replaced with fresh medium after 5 hours and the cells were imaged the next day.

siRNa treatment was performed using 2.5 *µ*L Lipofectamin^*TM*^ RNAimax (Ther-mofischer Scientific, # 13778075) and 1 *µ*L siRNA (20 *µ*mol.L-1 for a final concentration of 20 nmol.L-1) each mixed in 250 *µ*L OptiMEM for 5 min, then mixed and incubated at room temperature for 20 min. 300,000 cells were then resuspended, and plated in a 12-well plate in 500 *µ*L media total + siRNA mix. The cells were incubated with siRNA for 24h before experiments and qPCR. The RNA used were SMARTpool:ON-TARGETplus Pdcd6ip (Dharmacon #L-062173-01-0005) for Alix depletion and ON-TARGETplus Non-targeting Pool (Dharmacon #D-001810-10-05) as a scrambled control.

For live imaging of the spindle and post-mitotic bridges, tubulin was labeled using SIR-Tubulin (Tebu-bio #SC002, diluted in media to 20 nM and incubated for 6h then rinsed). These conditions were chosen because they allowed an optimal tubulin staining while not stabilizing the microtubules (as assessed by a normal duration of cell division).

### Cell sorting

Cells were sorted according to the fluorescence levels or forward scatter and side scatter to sort the cells by size (this recapitulates cell-cycle sorting (see Figure S2A) using FACSAria III Cell Sorter at the UCL flow cytometry core facility at UCL Great Ormond Street Institute of Child Health.

### Clonogenicity analysis

To test for speed and efficiency of exit from pluripotency, replating assays were performed. After various treatments such as sorting or culture on specific substrates, the cells were plated at low density (30,000 cells per well of a 24-well plate) onto plates coated with 0.1% gelatin in N2B27 for 26 hours. Then the cells were resuspended, counted, and replated at low density (200 cells per well of a 12-well plate) on 0.1% gelatin in N2B27+2i-LIF. After 5 days, the number of colonies was manually counted.

### Rex1-GFP intensity measurements in colonies and analysis

ES cells stably expressing Rex1-GFP and Gap43-mCherry (Kalkan et al., 2017) were plated in N2B27 on 0.1% gelatin-coated Ibidi dishes and 4 hours after plating, Z-stacks with ΔZ = 2 *µ*m were acquired. Rex1-GFP mean intensity was manually measured at the midplane of the cell using a rectangular selection for each cell at each time point using Fiji (42).

To determine whether the correlation between division and exit (Figure S1H) could be due to chance, we resorted to non-parametric bootstrapping methods. We first fitted the Rex1 intensity curves to a sigmoidal decay function and extracted the time of exit τ. We discarded the trajectories for which the time of exit τe could not be accurately determined (*i.e.* those where the fitting trajectories gave an error of fit for τe that was on the order of the value of τe itself). We then calculated the coefficient of determination R2 of the linear regression between the time of exit and the time of cell division (in case there were two events of division, the one closest to the timing of exit was picked). This gave R2=0.73 for the points that passed the criterion. We then bootstrapped the dataset by randomly assigning the time of exit of a cell i to the time(s) of cell division of a randomly selected cell j (the procedure was done with replacement) and calculated the coefficient of determination R2 for the randomized dataset. We performed this procedure 1000 times to build a distribution probability of correlation values. Importantly, we found that the observed correlation occurred just less than 5 of the time, underlying its statistical significance.

To determine the extent to which the dynamics of Rex1 downregulation in daughter cells were correlated (Figure 4E-F), we first separated cells exiting at the first division (where correlation between daughters was analyzed prior to them dividing again), and cell exiting at the second division (where correlation between grand-daughters was considered). In each case, we first calculated the average decay of Rex1 across all cells, as well as the standard deviation around it. This gave us a population-average of the variance (“global variance”) that would be observed if cells had no correlation from being sisters (to avoid artefacts due to differences in Rex1 expression levels between cells), we first normalized all Rex1 curves so that their first time point is of intensity 1). We then computed the variance at each time point between the Rex1 curves of two sisters (“local variance”). In both “global” and “local” case, we then averaged across time the variances, and compared the results. We only found two cases in which sister-sister variance was larger than the population average, which were the only two cases in which two sisters exited at different times. Importantly, looking at the full dataset, we found that the variance between sisters is typically 2-3-fold smaller than the global population variance, showing that sisters are heavily coordinated in their downregulation of Rex1.

### Volume measurements

Cell volumes were measured from Z-stacks using the 3D mesh plugin we previously published (22), https://github.com/PaluchLabUCL/DeformingMesh3D-plugin. The far-red membrane dye CellMask^*TM*^ (Thermofisher Scientific, C10046) was used for cell segmentation. The parameters used for segmentation were determined as the best by visual analysis. The parameters chosen were: gamma: 1000; alpha: 5; pressure: 0; normalize: 5; image weight: 1.0E-4; divisions: 3; curve weight: 0; beta: 0. The mesh deformation was made according to the perpendicular maximal gradient of the signal. The segmentation was stopped when the volume seemed resolved by visual assessment.

### Immunofluorescence

Cells were fixed in Ibidi dishes (IBI Scientific, 81156) in 4% formaldehyde in PHEM buffer with 0.125 % Triton. Primary antibodies against *α*-Tubulin (Thermo Fischer #62204 or Thermo Fischer Scientific #MA180017), and CRIK (Insight Biotechnology #611376) were incubated 1:200 in PBS with 5% non-fat dry Milk for 2h at room temperature, and the secondary antibody was incubated 1:500 for 1h at room temperature. Secondary antibodies were: Alexa Fluor® 647-AffiniPure Donkey Anti-Rat IgG (Stratech Scientific, #712-605-153-JIR), Goat anti-Rat IgG (H+L) Cross-Adsorbed Secondary Antibody, Alexa Fluor 568 (Thermo Fisher Scientific, #A-11077), Donkey anti-Mouse IgG (H+L) Highly Cross-Adsorbed Secondary Antibody, Alexa Fluor 488 (Thermo Fisher Scientific, #A-21202), Donkey anti-Rabbit IgG (H+L) Highly Cross-Adsorbed Secondary Antibody, Alexa Fluor 488 (Thermo Fisher Scientific, #A-21206), Donkey anti-Rabbit IgG (H+L) Highly Cross-Adsorbed Secondary Antibody, Alexa Fluor 647 (Thermo Fisher Scientific, #A-31573), Donkey anti-Rat IgG (H+L) Highly Cross-Adsorbed Secondary Antibody, Alexa Fluor 488 (Thermo Fisher Scientific, # A-21208), Donkey anti-Mouse IgG (H+L) Highly Cross-Adsorbed Secondary Antibody, Alexa Fluor 568 (Thermo Fisher Scientific, #A10037), Donkey anti-Rabbit IgG (H+L) Highly Cross-Adsorbed Secondary Antibody, Alexa Fluor 568 (Thermo Fisher Scientific, #A10042), Donkey anti-Rabbit IgG (H+L) Secondary Antibody, Alexa Fluor® 647 conjugate (Thermo Fisher Scientific, #A31573), Donkey anti-Rabbit IgG (H+L) Secondary Antibody, Alexa Fluor® 647 conjugate (Thermo Fisher Scientific, #A31573), Goat anti-Mouse IgG (H+L) Highly Cross-Adsorbed Secondary Antibody, Alexa Fluor Plus 647 ((Thermo Fisher Scientific, #A32728). The cells were mounted using ProLong® Gold Antifade Mountant with DAPI (Thermofisher Scientific, #P36941) and imaged using a 63X HCX PL APO (Numerical Aperture 0.6 - 1.4) on a confocal microscope (Leica DMI6000 Microscope).

### Substrate coating with E-Cadherin and laminin

Ibidi dishes (IBI Scientific, #81156) were plasma activated for 30 seconds and incubated overnight with 50 *µ*g.mL-1 E-Cadherin (RD Systems, #8875-EC-050) or 0.1% gelatin at room temperature, or 10 *µ*g.mL-1 Laminin (Sigma, #11243217001) at 37C.

### Microchannel experiments

PDMS microchannels were fabricated as previously described (43). Briefly, PDMS was polymerized over wafers containing 8 *µ*m*5 *µ*m or 10 *µ*m* 10 *µ*m channels and over 35 mm coverslips and pre-baked at 60C. Holes were punched on top of the channels then all the parts were attached to a dish and baked overnight at 60C. The channels were filled with media using syringes and let to equilibrate for 1h at 37 C and all bubbles were removed by softly pushing the gel down before the cells were injected using syringes.

For cells in microchannels, Rex1-GFP mean intensity was manually measured on the midplane of the cell every hour using a rectangular selection for each cell using Fiji (42).

### Single cell RNA sequencing

RNA Sequencing and Analysis: Library preparation was done by the Stem Cell Institute Genomics Facility using SmartSeq2 method using Nextera XT kits (Illumina) (44). Paired-end sequencing was performed on Illumina HiSeq4000 yielding 380 Million reads per lane.

#### RNA data processing and transcriptome analysis

Mouse genome build GRCm38/mm10 was used to align reads using GSNAP version 2015-09-29 (45). Genes were annotated using Ensembl release 81 (46) and read counts were quantified using HTSeq (47). Quality control and downstream analyses were performed using scran package in R (48). Expression was computed using DESeq2 package in R (49) with p-adjusted<0.05. Log-transformed normalized counts were used for subsequent heatmaps and expression plots. The cyclone function of the scran package was used to assign a cell cycle phase to individual cells (50).

#### Clustering of cells based on expression of two pluripotency genes Tfcp2l1 *Tbx3*

The two genes *Tfcp2l1* and *Tbx3* were selected because they are among the first genes to be downregulated upon exit from pluripotency (17), and they had the highest variation among all naive pluripotency genes when compared to naïve cells. Cells were assigned to one of 4 clusters using the k-mean clustering method, which minimizes the sum of squares of distance of each point to its cluster center. The clusters were then identified as high or low expression of each gene. Clustering was computed on normalized expression using DESeq2 (49) on both naive and exit cells together.

### qPCR

RNA extraction was performed using the RNA Easy Qiagen kit according to the manufacturer instructions. The reverse transcription was performed using the High-Capacity cDNA Reverse Transcription Kit (Thermofischer Scientific #4368814). qPCR were performed using SsoAdvanced(tm) Universal SYBR® Green Supermix (BioRad, #172-5271), loading 2.3 *µ*g per lane. Primers were bought from Integrated DNA technologies.

### FRAP

Cells were transfected with EGFP the day before the experiment and plated on laminin. FRAP was performed using an Olympus FluoView FV1200 Confocal Laser Scanning Microscope with 70% 405 nm laser on an ROI comprising most of the cytoplasm of the cell using the laser light stimulation (SIM) scanner with a 60X objective (UPLSAPO60XS, Numerical Aperture 1.3). Images were acquired at full speed (every 1.1s). The mean intensity on a ROI of fixed size was measured on the images.

### Statistical analysis

Prism 7 (Graphpad software, Inc) was used for all statistical analysis. The D’Agostino Pearson test was used to test for the normal distribution of data. To compare means, a Student t-test, a Student t-test with Welch correction or a Mann-Whitney test were performed if the data was normal with the same standard deviation, normal but with different standard deviation or not normal, respectively. For contingency data, *χ*^2^-tests were performed. N: number of independent experiments, n=number of points (not stated on dot plots).

## Table legends

**Table S1. The expression of key regulators of abscission does not strongly change during exit from naïve pluripotency.** Table showing the changes in RNA levels of key regulators of abscission during pluripotency exit; RNA levels at 24h or 25h after inducing exit are compared to levels in naïve cells. Black, data analyzed from (Kalkan et al., 2017); grey: data analyzed from (Yang et al., 2019).

## Movie legends

**Supplementary Movie 1. Rex1-GFP downregulation occurring shortly after a cell division.** Time-lapse (spinning-disk confocal microscopy) of ES cells expressing Rex1-GFP (green) and Gap43-mCherry (red) cultured in N2B27, following one dividing cell where Rex1-GFP intensity decreases after the first division. One frame is shown every 1h, starting 1h before division. One Z plane is shown. Scale bar: 10 *µ*m.

**Supplementary Movie 2. Rex1-GFP downregulation occurring shortly after a second cell division.** Time-lapse (spinning-disk confocal microscopy) of ES cells expressing Rex1-GFP (green) and Gap43-mCherry (red) cultured in N2B27, following one dividing cell where Rex1-GFP intensity decreases after the second division. One frame is shown every 1h, starting 1h before the first division. One Z plane is shown. Scale bar: 10 *µ*m.

**Supplementary Movie 3. ES cell dividing at the center of a colony.** Time-lapse (spinning-disk confocal microscopy) of a H2B-RFP (red) expressing naïve mESC colony labeled with CellMask^*TM*^ far red (cyan) with one cell dividing symmetrically at the center of the colony. One frame is shown every 5 minutes. One Z plane is shown. Scale bar: 10 *µ*m.

**Supplementary Movie 4. ES cell dividing at the border of a colony.** Time-lapse (spinning-disk confocal microscopy) of a H2B-RFP (red) expressing naïve mESC colony stained with CellMask^*TM*^ far red (cyan) with one cell dividing asymmetrically at the periphery of the colony. One frame is shown every 5 minutes. One Z plane is shown. Scale bar: 10 *µ*m.

**Supplementary Movie 5. HeLa cell division.** Time-lapse (spinning-disk confocal microscopy) of a S-HeLa colony labeled with Hoechst (red) and CellMask^*TM*^ (cyan). One picture is shown every 5 minutes. 0 min corresponds to NEBD. One Z plane is shown. Scale bar: 10 *µ*m.

**Supplementary Movie 6. Shape instabilities during ES cell divisions at the periphery of a colony.** Time-lapse (spinning-disk confocal microscopy) of a colony of naïve ES cells expressing H2B-RFP (red) and labeled with CellMask^*TM*^ far red (cyan) showing shape instabilities in cells dividing at the periphery of the colony. One frame is shown every 5 minutes. One Z plane is shown. Scale bar: 10 *µ*m.

**Supplementary Movie 7. Example of blebbistatin-treated ES cell dividing at the periphery of a colony.** Time-lapse (spinning-disk confocal microscopy) of a colony of naïve ES cells expressing H2B-RFP (red) colony and labelled with CellMask^*TM*^ far red (cyan) and treated with 1 *µ*M Blebbistatin with one cell dividing at the border of the colony. The cell displays strong size asymmetry between daughter cells. One frame is shown every 5 minutes. One Z plane is shown. Scale bar: 10 *µ*m.

**Supplementary Movie 8. Example of an ES cell dividing at the periphery of a 2D colony plated on E-Cadherin.** Time-lapse (spinning-disk confocal microscopy) of a naïve colony of ES cells expressing H2B-RFP (red) plated on E-Cadherin and labelled with CellMask^*TM*^ far red (cyan), with one cell dividing at the periphery of the colony. Cell division appears symmetric. One frame is shown every 5 minutes. One Z plane is shown. Scale bar: 10 *µ*m.

**Supplementary Movie 9. Example of an ES cell dividing at the periphery of a 2D colony plated on laminin.** Time-lapse (spinning-disk confocal microscopy) images of a cCellMask^*TM*^ aïve ES cells expressing H2B-RFP (red) and labelled with CellMaskTM far red (cyan), plated on laminin (see Methods section) with one cell dividing at the border of the colony. One frame is shown every 5 minutes. One Z plane is shown. Scale bar: 10 *µ*m.

**Supplementary Movie 10. ES cell dividing in confinement in a microchannels.** Time-lapse (spinning-disk confocal microscopy) of a H2B-RFP (red) expressing naïve ES cell dividing inside an 8*µ*m*5*µ*m PDMS microchannel (see Methods). The cell is stained with CellMask^*TM*^ far red (cyan) to highlight the membrane. One frame is shown every 5 minutes. The movie starts at metaphase. One Z plane is shown. Scale bar: 10 *µ*m.

**Supplementary Movie 11: Example of FRAP experiment highlighting exchange of cytoplasmic material between sister cells connected with a tubulin bridge.** Time-lapse (spinning-disk confocal microscopy) of ES cells expressing EGFP (fire LUT). The two upper cells are connected by a tubulin bridge. One frame is shown 11.1s before bleach, then one frame is shown every 1.1s. One Z plane is shown. Scale bar: 10 *µ*m.

**Supplementary Movie 12. Tubulin bridges in naïve ES cells.** Time-lapse (spinning-disk confocal microscopy) of a colony of naïve ES cells expressing H2B-RFP and labeled with SIR-Tubulin. One frame is shown every 15 minutes. A Z-projection over the entire volume of the colony is shown. Scale bar: 10 *µ*m.

**Supplementary Movie 13. Tubulin bridges in cells exiting naïve pluripotency (25 hours).** Time-lapse (spinning-disk confocal microscopy) of a colony of naïve ES cells expressing H2B-RFP after 25h of exit from pluripotency labelled with SIR-Tubulin. One frame is shown every 15 minutes. A Z-projection over the entire volume of the colony is shown. Scale bar: 10 *µ*m.

**Supplementary Movie 14. Tubulin bridge in cells exiting naïve pluripotency (45 hours)**. Time-lapse (spinning-disk confocal microscopy) of a colony of naïve ES cells expressing H2B-RFP after 45h of exit from pluripotency labelled with SIR-Tubulin. One frame is shown every 15 minutes. A Z-projection over the entire volume of the colony is shown. Scale bar: 10 *µ*m.

---

**Figure S1.**
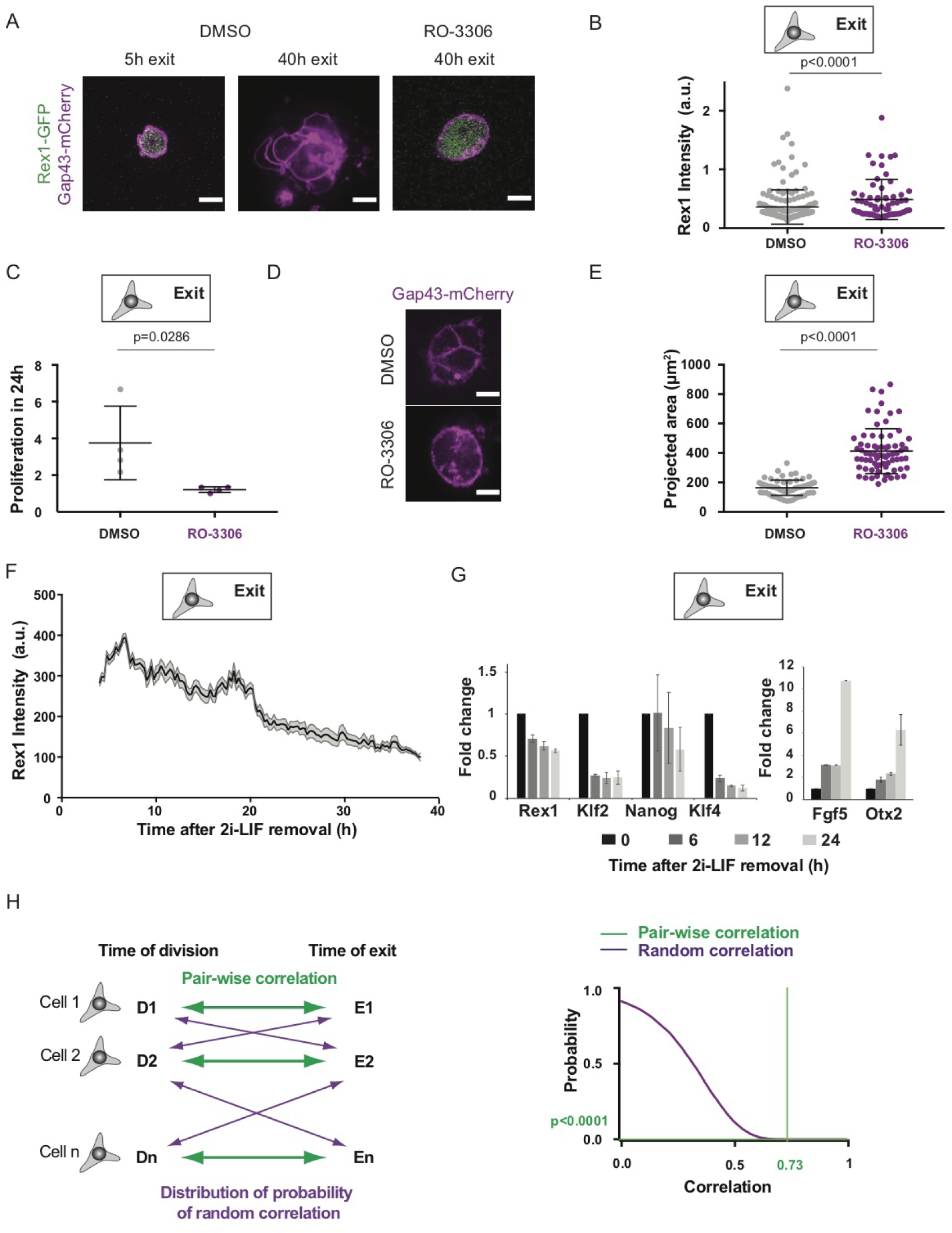
ES cells exit naïve pluripotency after mitosis (Related to Figure 1). A) Representative confocal microscopy images of ES cells expressing Rex1-GFP (green) and Gap43-mCherry (magenta), 5h or 40h after being placed in N2B27 supplemented with 6 *µ*M of RO-3306 or equivalent DMSO volume. Scale bars: 10 *µ*m. B) Dot plot showing Rex1-GFP intensity 40h after removal of 2i-LIF for cells treated with 6 *µ*M RO-3306 (purple) or DMSO (control, grey). The mean and standard deviation are shown. N=2. C) Dot plot showing proliferation rate over 24h of ES cells treated with DMSO (grey) or 6 *µ*M RO-3306 (purple). The mean and standard error of the mean are shown. N=5. D) Representative confocal images showing cell areas 40h after 2i-LIF removal and treatment with DMSO (top) or 6 *µ*M RO-3306 (bottom). Gap43-mCherry (magenta) is shown as a membrane marker. One Z-plane is shown in the middle plane of the cell. E) Dot plot showing the area of ES cells 40h after removal of 2i-LIF in presence of DMSO (grey) or 6 *µ*M RO-3306 (purple). The mean and standard deviation are shown. N=2. F) Plot of Rex1-GFP mean intensity in ES cells expressing Rex1-GFP and Gap43-mCherry during exit from pluripotency as a function of time. 0h: time of 2i-LIF removal. The mean and standard error of the mean are plotted. N=3, n= 33. G) Bar graphs showing the expression of key pluripotency genes (left) or exit genes (right), as assessed by qPCR during exit from pluripotency. The mean and standard error of the mean are shown. N=3. H) Left: schematic outlining the analysis of the robustness of the correlation between time of exit and time of division. Right: cumulative probability of correlation coefficients R2 (see Methods for details) arising at random (purple), plotted together with the experimental correlation (green).

**Figure S2.**
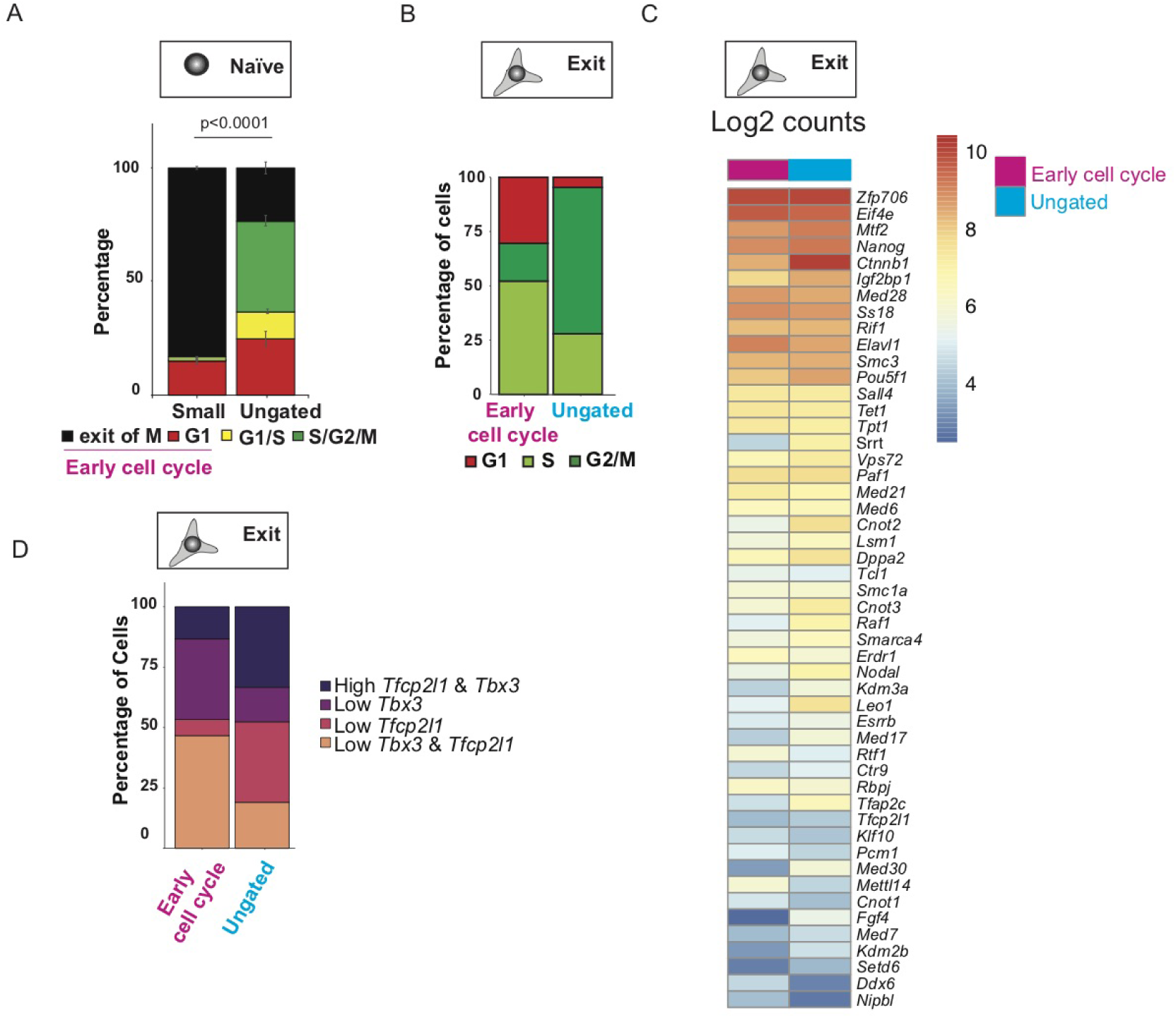
Inducing pluripotency exit early in the cell cycle leads to faster loss of naïve pluripotency (Related to Figure 2). A) Bar graph showing the percentage of cells in each phase of the cell cycle assessed by comparing the Fucci2a levels in the cell sorting by cell size experiment; the “small” cell population is compared to the ungated population (G1: red; G1/S: yellow; S/G2/M: green; black: exit of mitosis). The mean and standard error of the mean is plotted. N=2. B) Bar graph showing the percentage of cells in each phase of the cell cycle, assessed by sc-RNASeq analysis, 6h after triggering pluripotency exit in cell populations sorted by size (the “early cell cycle” population corresponds to the cells sorted as small, see (A)). (G1: red; S: light green; G2/M: green). C) Heat-map showing the Log2 counts of the levels of expression of the main pluripotency genes for the two cell populations obtained by sorting cells by size, and placed in N2B27 for 6h. Early cell cycle (small cells): pink; ungated: blue. D) Bar graph showing the percentage of cells displaying high or low expression levels of *Tfcp2l1* and *Tbx3* as a proxy for levels of naïve pluripotency (see Methods). Early cell cycle: pink; ungated: blue.

**Figure S3.**
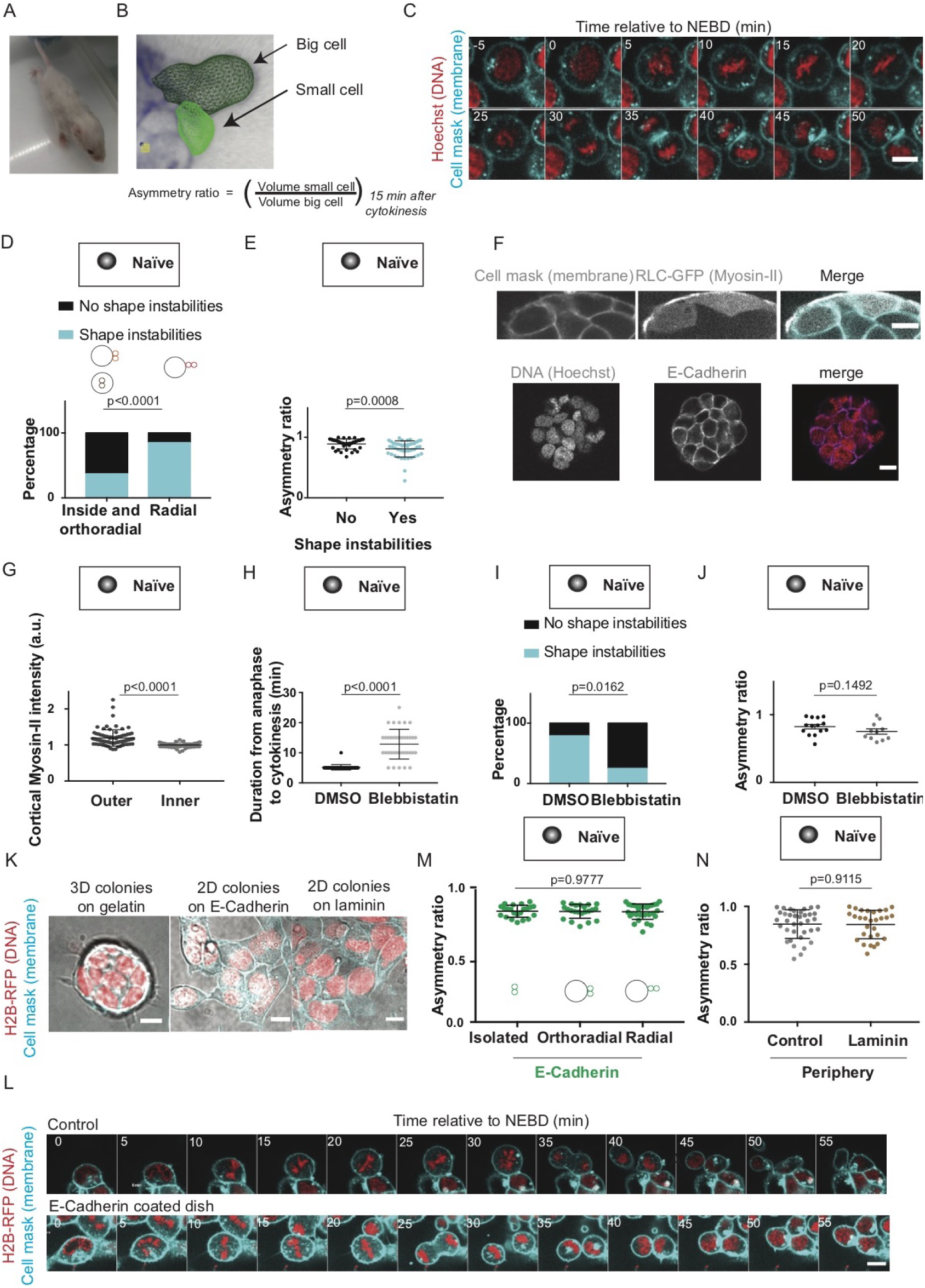
Mechanisms controlling division asymmetries in ES cells (Related to Figure 3). A) Picture of a chimera mouse obtained by injection of the H2B-RFP cells into a C57B6 blastocyst. B) Image: 3D rendition of 2 daughter cells labeled with CellMaskTM far red and segmented using the 3D mesh plugin (see Methods). The arrows indicate the small and big sister cells. Bottom: definition of the asymmetry ratio. C) Time-lapse spinning-disk confocal microscopy images of a S-HeLa labeled with Hoechst (red) and CellMaskTM far red (cyan). Time in min; 0 min: time of nuclear envelope breakdown (NEBD). One Z-plane is shown. Scale bar: 10 *µ*m. D) Bar graph showing the percentage of cells displaying shape instabilities (blue) or no shape instabilities (black) as assessed visually for naïve H2B-RFP ES cells dividing inside or at the periphery of a colony with the spindle parallel to the colony border (left), or at the periphery of a colony with the spindle perpendicular to the colony border (right). N=3, n= 84. E) Dot plot showing the asymmetry ratio between sister cells for naïve H2B-RFP ES cells displaying (right) or not displaying (left) shape instabilities. The mean and standard deviation are shown. N=3. F) Top: confocal images of naïve ES cells expressing H2B-RFP, RLC-GFP and labeled with CellMask^*TM*^ far red. Bottom: confocal images of naïve ES cells stained for E-Cadherin and DNA. Scale bars, 10 *µ*m. G) Dot plot showing cortical Myosin-II intensity in naïve ES cells expressing RLC-GFP. The mean intensity as measured manually of the outer cortex (left) and inner cortex (right) and standard deviation are shown. N=2. H) Dot plot showing the time between anaphase and cytokinesis in ES cells treated with DMSO (black) and with 1 *µ*M Blebbistatin (grey). The mean and standard deviations are shown. N=2. I) Bar graph showing the percentage of cells displaying shape instabilities (blue) or no shape instabilities (black) for naïve ES cells dividing at the periphery of a colony treated with DMSO (left) or 1 *µ*M Blebbistatin (right). The percentage is shown. N=2, n=14 for DMSO, n=12 for Blebbistatin. J) Dot plot showing the asymmetry ratio for ES cells treated with DMSO and 1 *µ*M Blebbistatin. The mean and standard deviation are shown. N=2. K) Representative images of colonies of naïve ES cells expressing H2B-RFP (red) and labelled with CellMask^*TM*^ far red (cyan) on a gelatin-coated substrate (left), an E-Cadherin-coated substrate (middle) or a laminin-coated substrate (right). A Z-projection overlaid on a transmitted light image is shown. Scale bar: 10 *µ*m. L) Representative time-lapse of dividing naïve ES cells expressing H2B-RFP (red) and labeled with CellMask^*TM*^ far red (cyan) on a gelatin-coated substrate (top) or an E-Cadherin-coated substrate (bottom). Time in min; 0 min: time of NEBD. One Z-plane is shown. Scale bar: 10 *µ*m. M) Dot plot representing the size asymmetry ratio between daughter cells at division for naïve ES cells plated on E-Cadherin and dividing as single cells (“isolated”) or at the periphery of a colony with the spindle oriented orthoradially or radially to the colony border. The mean and standard deviation is plotted. N=2. N) Dot plot showing the asymmetry ratio for naïve ES cells dividing on gelatin (control) or laminin. The mean and standard deviation are shown. N=3.

**Figure S4.**
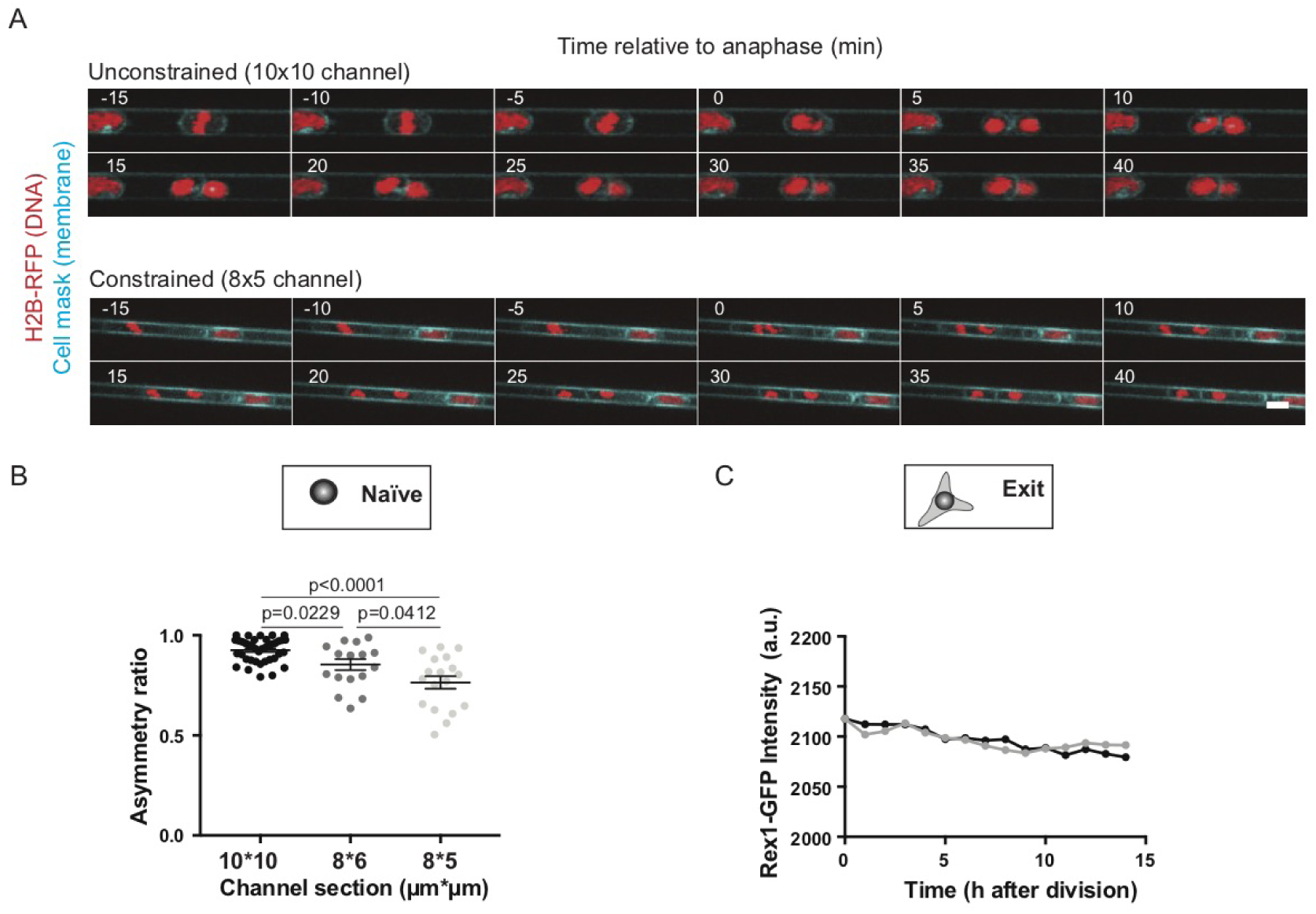
Confinement induces strongly asymmetric divisions with no effect on exit from naïve pluripotency dynamics (Related to Figure 3). A) Time-lapse spinning-disk confocal microscopy images of a naïve ES cell expressing H2B-RFP (red) and labeled with CellMask^*TM*^ far red (cyan) dividing in a 10*10 *µ*m channel (top) or an 8*5 *µ*m channel (bottom). One Time in min; 0 min: anaphase. One Z-plane is shown. Scale bar: 10 *µ*m. B) Dot plot representing the sister cell asymmetry ratio of H2B-RFP ES cells dividing in 8*5 *µ*m, 6*8 *µ*m or 10*10 *µ*m channels. The mean and standard deviation are plotted. N=2 for each condition. C) Graph showing an example of Rex1 intensity time course for 2 daughter cells resulting from an asymmetric division in a 8*5 *µ*m microchannel. 0h: time of division.

**Figure S5.**
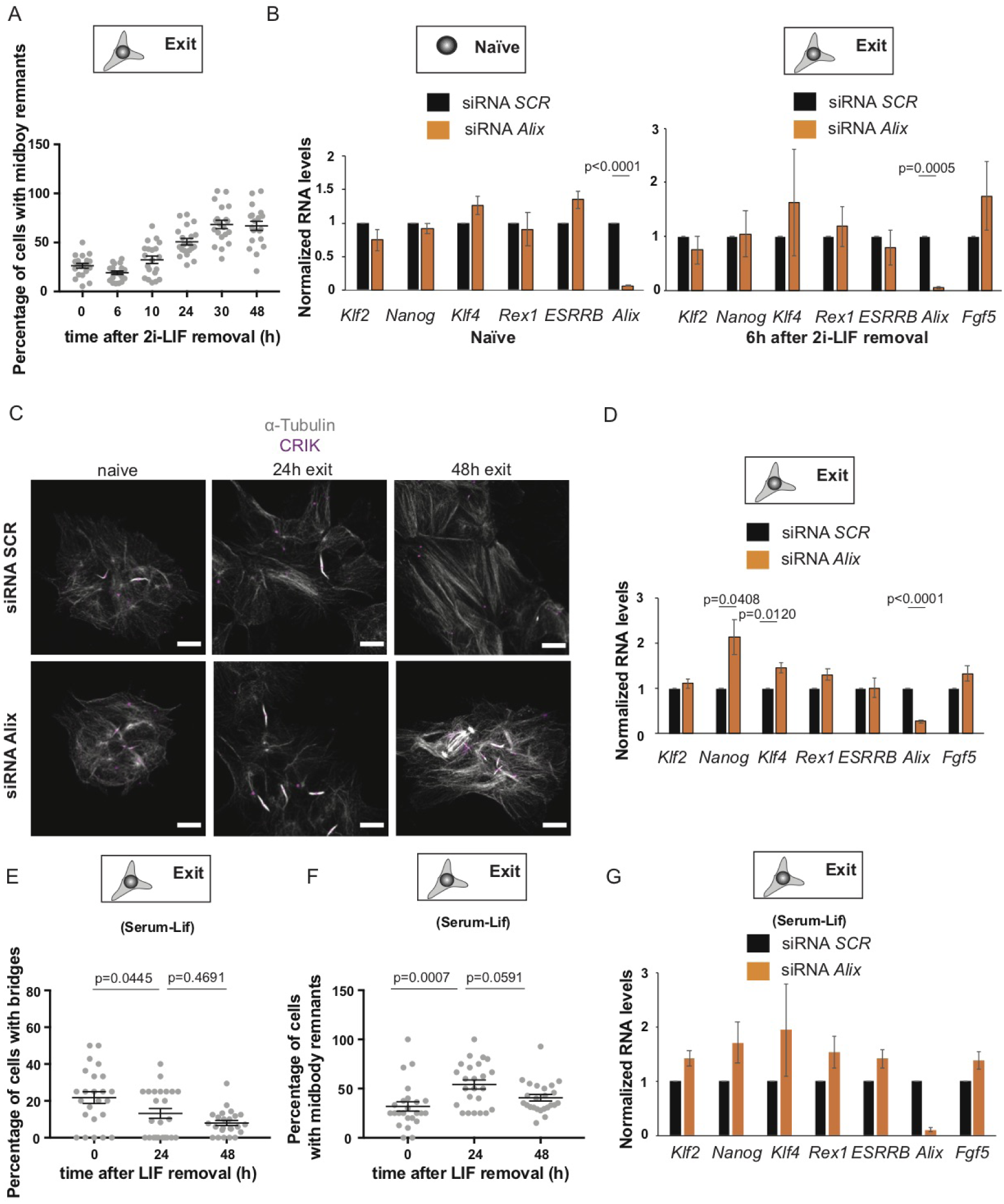
Interfering with abscission impairs exit from naïve pluripotency (Related to Figure 5). A) Dot plot showing the percentage of cells associated with a midbody remnant (*i.e.* CRIK spot not attached to a bridge) detected in colonies of naïve H2B-RFP ES cells and during exit from pluripotency. The mean and standard error of the mean are shown. N=2. B) Box plot showing RNA levels for H2B-RFP mESC ES cells treated with Scrambled siRNA (SCR, black) or siRNA against Alix (orange) for 24h in 2i-LIF (left) or for 24h in 2i-LIF followed by 6h in N2B27. The mean and standard error of the mean are shown. N=5 for 2i-LIF and N=2 for 6h exit. C) Confocal image of H2B-RFP ES cells treated with Scrambled siRNA (SCR, top) or siRNA against Alix (bottom) for 24h in 2i-LIF then plated on laminin in 2i-LIF or N2B27 and stained for *α*-Tubulin (white) and CRIK (magenta). A maximum Z-projection over the volume of the colony is shown. Scale bars: 10 m. D) Box plot showing RNA levels for H2B-RFP ES cells treated with Scrambled siRNA (SCR, black) or siRNA against Alix (orange) for 24h in 2i-LIF followed by 24h in N2B27. The mean and standard error of the mean are shown. N=5. E) Dot plot showing the percentage of cells with bridges in naïve H2B-RFP ES cell colonies maintained in Serum-LIF or allowed to exit from pluripotency for 24 or 48h. The mean and standard error of the mean are shown. N=2. F) Dot plot showing the percentage of cells associated with midbody remnants (*i.e.* not attached to a bridge) per cell detected in naïve H2B-RFP ES colonies maintained in Serum-LIF or allowed to exit from pluripotency for 24 or 48h. The mean and standard error of the mean are shown. N=2. G) Box plot showing RNA levels for H2B-RFP ES cells treated with Scrambled siRNA (SCR, black) or siRNA against Alix (orange) for 24h in Serum-LIF then cultured in N2B27 for 24h. The mean and standard error of the mean are shown. N=2.

